# 5’XP sRNA-seq: Efficient Identification of Transcripts With and Without 5’ Phosphorylation Reveals Evolutionary Conserved Small RNA

**DOI:** 10.1101/2020.08.21.261412

**Authors:** Unn Kugelberg, Daniel Nätt, Signe Skog, Claudia Kutter, Anita Öst

## Abstract

Small RNA (sRNA) sequencing has been critical for our understanding of many cellular processes, including gene regulation. Nonetheless, the varying biochemical properties of sRNA, such as 5’ nucleotide modifications, makes many sRNA subspecies incompatible with common protocols for sRNA sequencing. Here we describe 5XP-seq that outlines a novel strategy that solves this problem. By tagging 5’P sRNA during library preparation, 5XP-seq combines an open approach that includes all types of 5’-terminal modifications (5’X), with a selective approach for 5-phosphorylated sRNA (5’P). We show that 5XP-seq not only enriches phosphorylated miRNA and piRNA but successfully discriminates these sRNA from all other sRNA species. We further demonstrate the importance of this strategy by successful inter-species validation of sRNAs that would have otherwise failed, including human to insect translation of several tRNA (tRFs) and rRNA (rRFs) fragments. By combining 5’ insensitive library strategies with 5’ sensitive tagging, we have solved an intrinsic bias in modern sRNA sequencing that will help us reveal the true complexity and the evolutionary significance of the sRNA world.

## BACKGROUND

High-throughput RNA-sequencing techniques have revolutionized our understanding of various small noncoding RNA (sRNA) species, typically with an arbitrarily defined length of less than 200 nucleotides. Numerous studies have provided insight into the indispensable functions of sRNAs in regulating cellular processes, such as cell development, differentiation and proliferation under physiologically normal and pathological conditions [1, 2]. In addition to well-characterized micro (miRNA), small interfering (siRNA) and PIWI-interacting RNAs (piRNA)[3, 4], new sRNA species deriving from transfer (tRNA), ribosomal (rRNA) [5], small nucleolar (snoRNA), vault (V RNA) and Y RNA [6] exists in many cell types including the transcriptionally dormant sperm [7]. These fragments are not just by-products of pervasive transcription or random RNA degradation but are precisely engineered to exert specialized regulatory functions. For example, specific fragments generated from tRNAs (tsRNA) have been implicated in translational control and cell growth [8]. The regulatory repertoire of sRNA in general, and tsRNA specifically, is expanded by the addition of small molecular modifications including methylations and pseudouridylation. Such modifications effect RNA stability, three-dimensional structure, cellular location and interactions with proteins [9-11].

A major limitation when exploring the sRNA world is the biased enrichment of different sRNA species introduced during library preparation that are caused by diverse chemical modifications at the 5’- and 3’-ends of RNA. Hydroxyl (OH) or phosphate (P) groups can be found either in the 3’- or 5’-end depending on the processing pathway, while a 2’,3’-cyclic phosphate (cP) group is only present on the 3’-end [12]. In addition, sRNAs can have multiple types of 5’-caps such as mGpppC, 7mGpppG, GpppG, GpppA, and 7mGpppA in human cells [13] or NAD+ [14] and 3′-dephospho-CoA, succinyl-dephospho-CoA and acetyl-dephospho-CoA in bacteria [15].

Most high-throughput sequencing methods require the ligation of sequencing-compatible adapters first to a hydroxyl-group at the 3’-end (3’-OH) and then to a phosphate group at the 5’-end (5’-P) of the sRNA molecule. These chemical reactions therefore select for sRNA containing these very specific terminal modifications, that are subsequently PCR amplified and finally detected during sequencing.

Here, we describe 5’XP sRNA-seq, a novel sequencing method with a unique tagging system that allows the detection of both sRNA with a 5’-P (typically miRNA and piRNA) and sRNA with alternative 5’ terminal groups. We show that this method increases specificity towards miRNA and piRNA when compared to a commonly used strategy, while simultaneously allowing for additional fragments to be detected in the same library. Overall, we show that 5’XP sRNA-seq is an adequate strategy for tackling critical biases commonly prevalent in standard next generation sRNA library preparations.

## RESULTS

### 5’P, 5’X and 5’XP sRNA-seq employ distinct library preparation strategies

5’-P and 3’-OH dependent ligations amplify some sRNA species, including piRNA and miRNA, but will exclude functional sRNA species without a 5’-P. We refer to this ligation strategy as 5’-P dependent sRNA-seq (from here on: 5P-seq). This should not be mistaken for 5’-P dependent methods recently developed for long RNA sequencing [16-18]. It has been shown that the 5’P-ligation is sensitive to secondary structures of the RNA, which further restricts capturing a broad range of sRNA species (for review see [19]). To include highly structured RNA species without 5’-P ends, strategies have been developed in which the 5’ adapter is not ligated to the RNA but is added later to the cDNA [20, 21]. We here refer to such methods as 5’ inclusive sRNA-seq (5X-seq). Building on these two approaches we developed 5’XP sRNA-seq (5XP-seq). In this method, the first adapter is ligated to the 3’ end and a 11-nt long oligo is ligated to the 5’ end of sRNA, which allows efficient tagging of RNAs with 5’-P. Then, after reverse transcription of the RNA ligation product, a sequencing-compatible adapter is ligated to the corresponding 3’ end of all cDNA (Fig 1, Supplementary Fig. 1). Through this approach sRNA with 5’-P and 5’-X can be sequenced in the same library and then bioinformatically discerned based on the presence or absence of the 5’ oligonucleotide tag.

**Fig 1.**
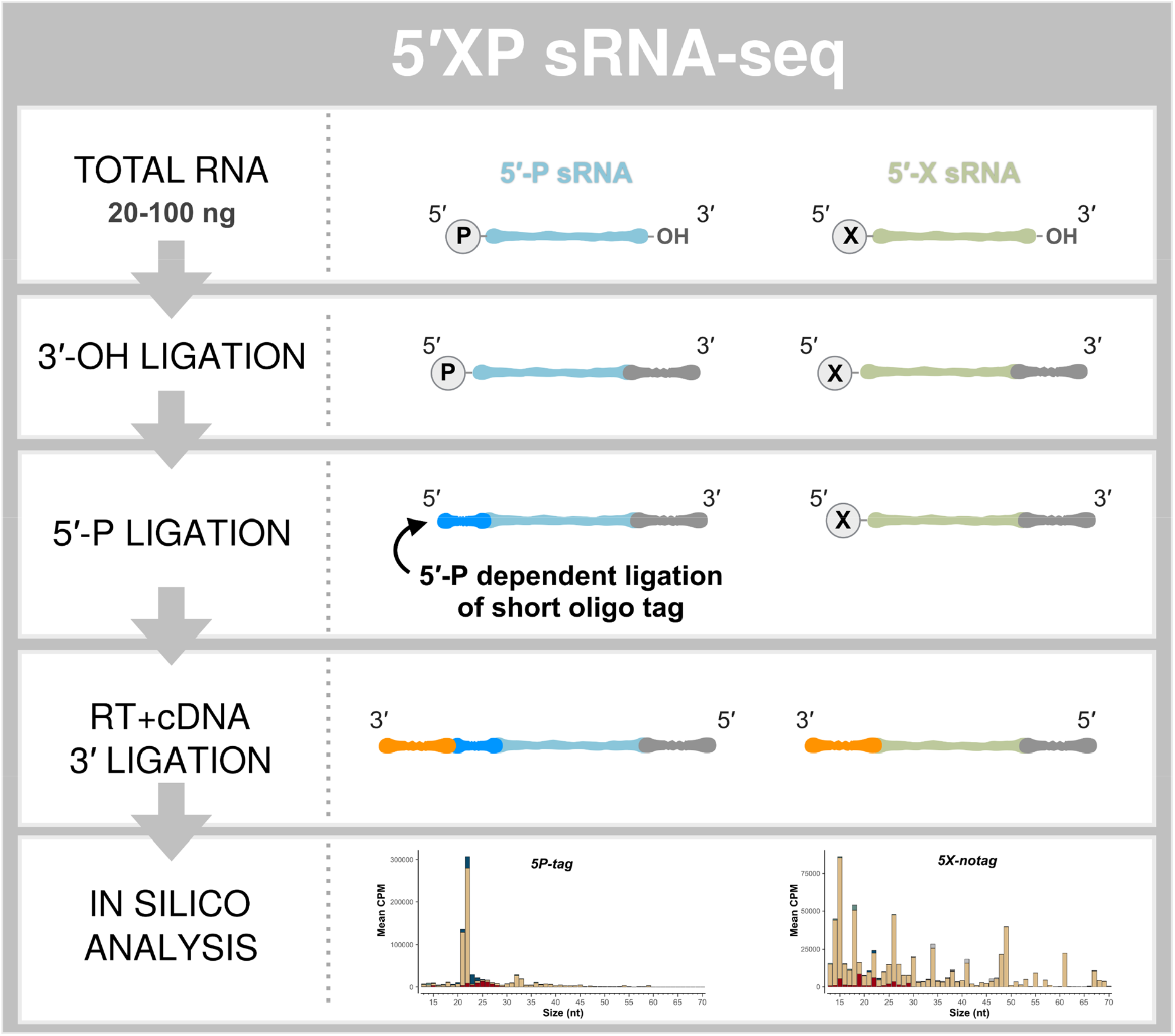
The principle steps of 5′XP sRNA-seq. Small RNA sequencing using 5′XP sRNA-seq generates two sub-libraries, either sensitive (5P-tag) or insensitive (5X-notag) to a phosphate (P) in the 5′-terminal of the original RNA. This is done by tagging RNA fragments with 5′-P using a sequence specific oligo that identifies the 5′-P in the downstream bioinformatic analysis.

### 5P and 5X sRNA-seq generate libraries with substantially different sRNA content

To better understand how 5P and 5X sRNA-seq vary in their enrichment for different RNA species, we first compared libraries generated using a popular commercial 5P-seq kit (NEBNext Multiplex Small RNA Library prep kit for Illumina; New England Biolabs) and an adaptation of published 5X-seq protocols [20-23]. For comparative reasons, we modified the 5X-seq protocol to work with the reverse transcriptase of the 5P-seq kit. We extracted total RNA from a pool of 50 *Drosophila* embryos, 0.5-2.5 hours old, which was used as input for all sequencing library preparations. Each preparation used only 30 ng of total RNA that equals to one third of the RNA extracted from one single drosophila embryo, rendering the protocols highly efficient. We adopted sequence-based counting of the reads, for which statistical units are based on counts per sequence and not counts per feature (see methods for details). An annotated count table normalized against total library sizes as counts per million (cpm) are presented in Supplementary Table 1. Raw data is available at Sequence Read Archive under the accession number PRJNA658107.

**Table 1.**
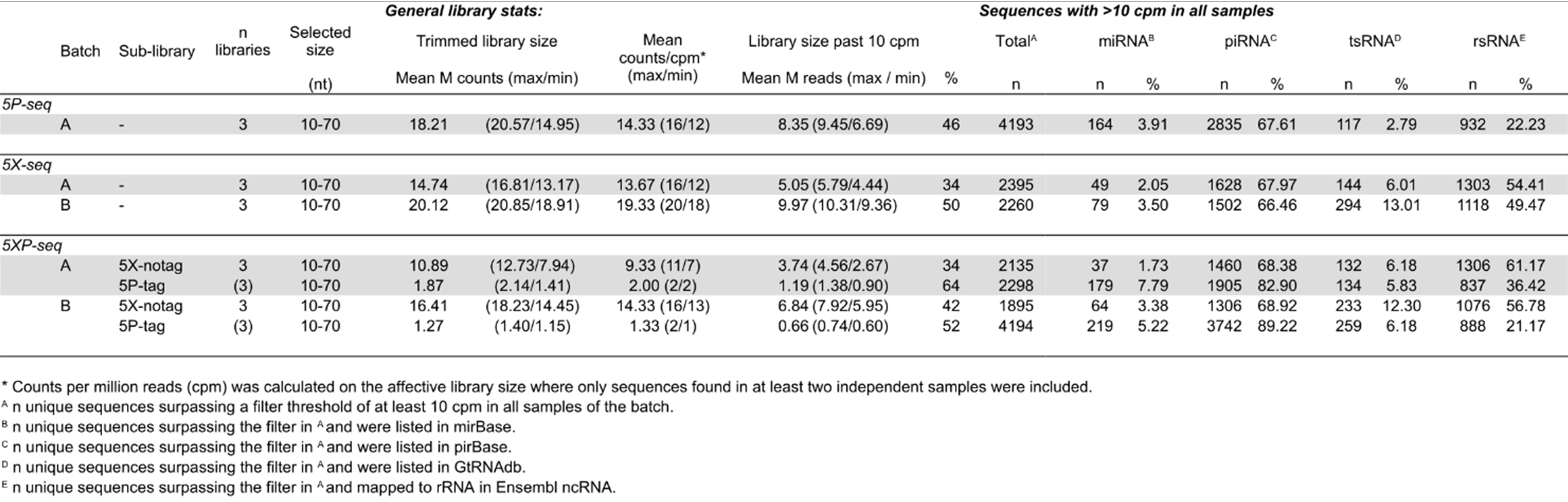
Library stats and sequence diversity

Library stats for the 5P- and 5X-type libraries using single end sequencing for 75 cycles are found in Table 1. Data from 5P-seq libraries were in accordance with our previous observations using this kit. To better validate the 5X-seq protocol, we generated a second batch of 5X-seq using independent embryos (Batch B). Focusing on intermediately-to-strongly expressed sRNA (>10 cpm), 5P-seq resulted in more alignments against miRNA and piRNA, and less alignments against rRNA and tRNA, when compared to 5X-seq (Fig 2). This result was similar for both cumulative counts (Fig 2A) and relative fold changes (Fig 2B), indicating a general trend across most sRNA. This difference is expected since miRNA and piRNA are commonly 5’ phosphorylated and should be preferably enriched by 5P-seq, while tRNA and rRNA show more diversity in the 5’-end and should preferably be caught by 5X-seq. When not allowing mismatches in the annotation 5X-seq annotated better to the references than 5P-seq (light grey Fig 2A-B). The advantage for applying sequence-based counting is that mismatches in the alignments between sequenced reads and small RNA reference sequences only affect the annotation without changing the raw count table, which is often the case for feature-based counting. Allowing up to 3 mismatches when annotating sequences against small RNA references removed nearly all sequences with no annotation in 5P-seq (light grey Fig 2C-D). Since libraries were generated from the same RNA pool, this indicates that 5P-seq may have suffered from misreading during cDNA-synthesis possibly by interference from RNA-modifications. Removing sequences mapping to rRNAs, however, also removed this difference (Supplementary Fig 2).

**Fig 2.**
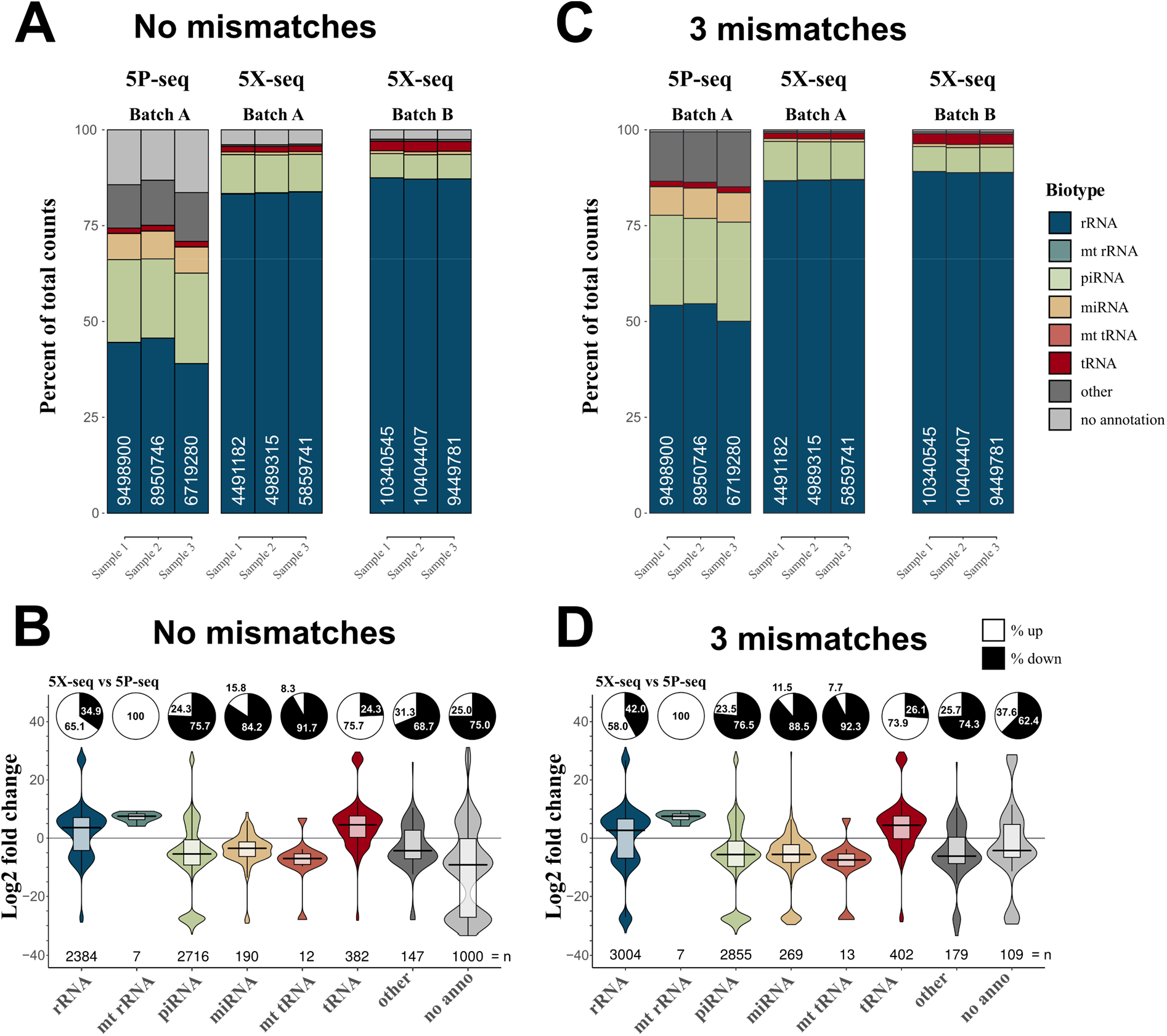
5X-seq catches different small RNAs compared to 5P-seq. In order to generate a sequencing method more inclusive of RNAs with 5′ modifications we compared two adapter ligation strategies: 5P-seq adds the 5′ adapter on RNA and is dependent on 5′ phosphorylation, and 5X-seq adds the 5′ adapter, to the corresponding 3′ side of cDNA and is therefore independent of 5′ RNA phosphorylation. Stacked bars showing the sRNA composition of 5P and 5X libraries, allowing for **(A)** no mismatches and **(C)** 3 mismatches in the annotation against sRNA reference sequences. Batch A and B indicates samples isolated from the two independent pools of RNA. Violin plots with log2 fold differences between 5X and 5P per unique sequence when **(B)** 0 mismatches and **(D)** 3 mismatches are allowed. Pie charts show percent up and down regulated sequences. Only unique sequences that passed 10 CPM in all samples of a method (either 5P or 5X) was included. mt=mitochondrial. CPM = counts per million reads.

While the majority of sRNA identified by 5X-seq were also present in 5P-seq libraries, some sRNAs were completely absent in one or the other library, which resulted in a more than 20-fold change (Fig 2B and 2D; Supplementary Table 1). This indicates substantial differences in the composition of sRNA species identified by the two different library strategies. Therefore, studies that have applied only 5P-based sRNA-seq strategies may have misinterpreted the proportions of small RNA present in their samples.

### 5XP sRNA-seq distinguishes piRNA and miRNA

To preserve sequence diversity given the observed biases described above, in parallel to the 5P and 5X-seq libraries, we generated 5XP-seq libraries using the same two pools of RNA from *Drosophila* embryos (Batch A and B). Table 1 summarizes the general properties of 5XP-seq after being sorted into the two sub-libraries, either containing (5P-tag) and not containing (5X-notag) the 5’ oligo tag. In theory, 5P-tag and 5X-notag of 5XP-seq would correspond to the 5P-seq and 5X-seq libraries, respectively.

Similar to the difference observed between 5P- and 5X-seq, 5XP-seq had more specificity towards miRNA and piRNA in the 5P-tag sub-libraries, than in the 5X-notag sub-libraries (Table 1). In fact, sRNA diversity was highly similar between 5X-notag sub-libraries and regular 5X libraries prepared without tagging 5’-P (Table 1). In support, expression levels of 5X-seq and 5X-notag cohered strongly, both by correlation and unsupervised clustering (Fig 3A).

**Fig 3.**
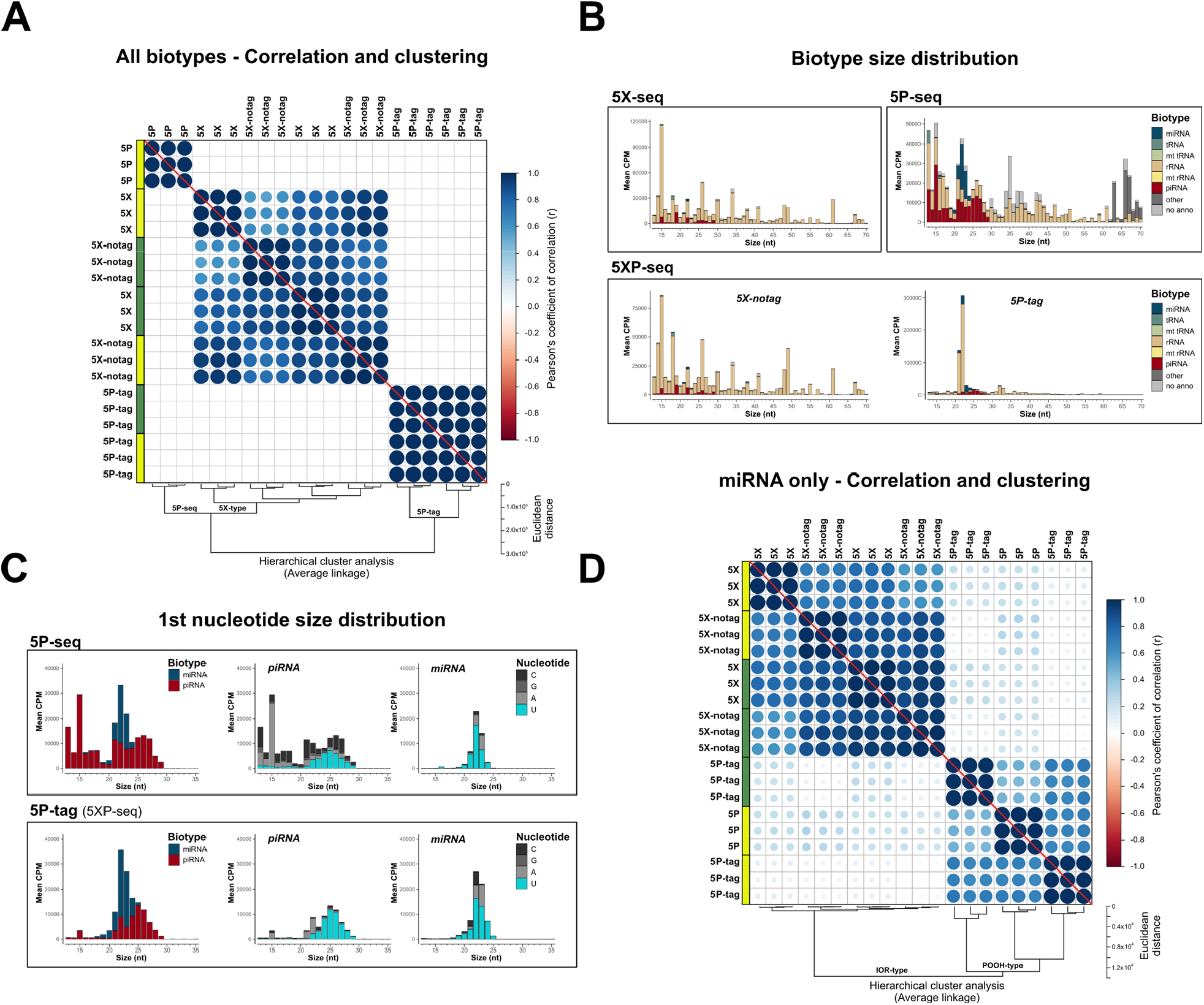
5XP-seq replicates 5X-seq and identifies 5′ phosphorylated sRNA species. 5XP-seq distinguishes sRNA with (5P-tag) and without (5X-notag) 5′-terminal phosphorylation which characterize canonical miRNA and piRNA. **(A)** Correlation plot ordered by unsupervised hierarchical clustering showing three separate clusters, where all 5X-type libraries (5X-seq; 5X-notag sub-libraries of 5XP-seq) generates similar sequencing libraries, while 5P-seq and the 5P-tag sub-libraries of 5XP-seq forms two separate clades. **(B)** Histograms showing the mean expression of biotypes across different fragment sizes. **(C)** Size distributions of known miRNA and piRNA (left panels) show that the 5P-tag sub-libraries of 5XP-seq generates a cleaner canonical piRNA peak in the expected size range, with a typical Uridine (U) bias on the 1st nucleotide (middle panels), than 5P-seq. Size distribution and 1st nucleotide bias of miRNA are more similar between libraries (right panel). **(D)** Correlation plot ordered by hierarchical clustering of only annotated miRNA confirms that 5P-seq and 5P-tag sub-libraries enrich the same miRNA. Analysis was performed on sRNA that reached 10 CPM in all replicates of a method. Yellow and green boxes in the correlation plots shows libraries generated from two separate pools of RNA from *drosophila* embryos. CPM = counts per million reads.

In contrast, we determined higher specificity (∼ 2-fold) of miRNA and piRNA species in 5P-tag sub-libraries when compared to regular 5P-seq libraries (Table 1). This indicates that ligating a short oligonucleotide tag at the 5’-terminal—as performed in 5XP-seq—captures 5’-P sRNAs more efficiently when compared to the long 5’ adapter ligation used in regular 5P-seq. Nonetheless, the 5P-tag sub-libraries only contributed with 10% to 20% of the total library sizes in 5XP-seq (Table 1) and correlated poorly with regular 5P-seq (Fig 3A). This led to two possible explanations. First, 5’-P sRNA may constitute a smaller proportion of the total sRNA population. Secondly, 5XP-seq may suffer from inefficient ligation of the 5’ oligo tag. In support of the former, a saturation analysis showed that the 5P-tag sub-libraries were equally saturated to the other libraries. Thus, irrespective of library-type, increasing the sequencing depths would result in little gain in identifying new sRNA specimens from the sRNA population targeted by each method (Supplementary Fig 3).

To further test the hypothesis that the 5P-tag sub-library of 5XP-seq targets a smaller and possibly more specific population of 5’-P sRNAs, we characterized each library type in more details. Size distribution of different sRNA species confirmed a sharp peak at 22 nt in 5P-tag sub-libraries, while the other library strategies showed more diversity (Fig 3B). The peak at 22nt is the typical size of miRNA. However, we detected that that many sequences of 22nt also mapped to rRNAs. Therefore, we extracted miRNA and piRNA that are known 5’-P subspecies for further comparison. Both the 5P-tag sub-libraries and regular 5P libraries where enriched with a sharp 22-23 nt miRNA peak, and a broader 22-29 nt piRNA peak (left panel Fig 3C). However, the 5P-tag sub-libraries of 5XP-seq had fewer non-canonical piRNA fragments that were shorter than 21 nt and showed a higher proportion of canonical piRNA with uridine bias at their first nucleotide (middle panel Fig 3C; [24]). Interestingly, the first nucleotide uridine bias was also observed in miRNA (right panel Fig 3C), which could indicate a new subspecies of miRNA in early drosophila embryos.

A combination of a lower proportion of unannotated small RNA (light grey in Fig 3B and 2A), as well as a smaller target population of sRNA (Supplementary Fig 3) and more canonical piRNA profiles (Fig 3C), altogether indicated that the 5P-tag sub-library of 5XP-seq enriches for 5’-P RNA better than regular 5P-seq. The low correlation in the expression profiles observed between these two strategies (Fig 3A) was also rescued (Fig 3D) after noise from less studied subspecies of sRNA was reduced by analyzing miRNAs only, which represent the most well-characterized 5’-P sRNA species.

### Library strategies are defined by opposite terminal enrichments of small rRNA

Loss of peak integrity of rRNA subunits is often used as an indicator of RNA degradation in studies of long RNA, often referred to as the RNA integrity number (RIN). While a similar measurement is lacking for sRNA, all our libraries showed very sharp peaks indicating specifically processed rRNA with high sRNA integrity (Fig 4A-B).

**Fig 4.**
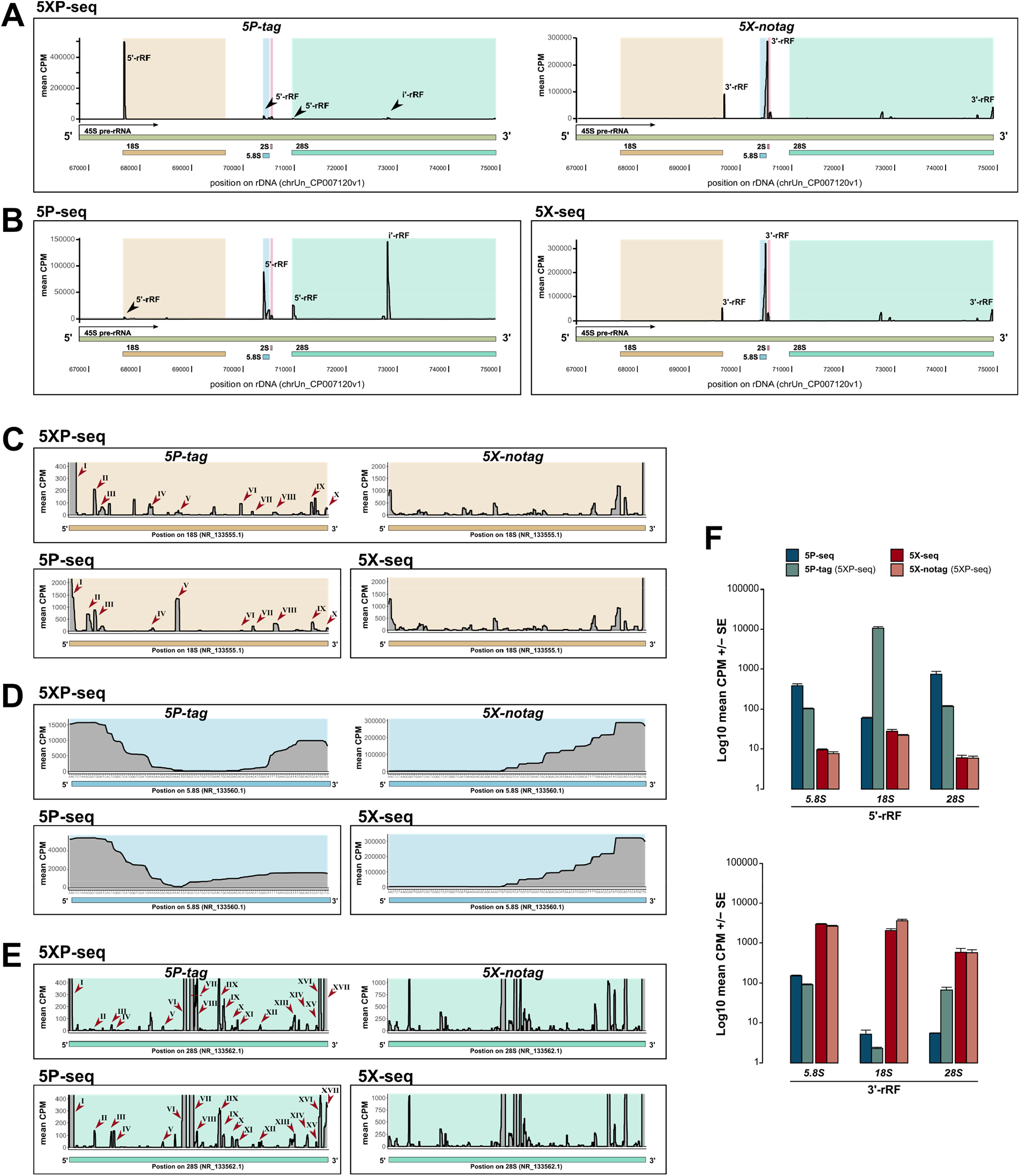
5P-type libraries enrich 5’ rRNA fragments while 5X-type libraries enrich 3’ fragments. Graphs shows 5’ and 3’ stratification in sRNA derived from rRNA (rRF). **(A-B)** Coverage plots of rRFs across an example pre-rRNA sequence (45S; green) located on the repetitive rDNA loci (chrUn_CP007120v1), annotated with mature rRNA subunits 18S (light brown), 5.8 (light blue), 2S (light red) and 28S (light turquoise). **(A)** Shows mean CPM coverage of 5XP-seq libraries separated by 5P-tag (left panel) and 5X-notag (right panel) sub-libraries. **(B)** Instead shows 5P-seq and 5X-seq control libraries. **(C-E)** Zoomed-in coverage plots of the 18S, 5.8S and 28S subunits. For 5P-type libraries, roman numbers indicate major peaks present in both regular 5P-seq libraries and the 5P-tag sub-libraries of 5XP-seq. **(F)** Bars show 5’-rRF enrichment in 5P-type libraries (upper panel) and 3’-rRF enrichment in 5X-type libraries (lower panel). CPM = counts per million reads.

The 5P-tag sub-libraries in 5XP-seq were dominated by a peak centered at 22 nt (Fig 3B), which primarily contained an rRNA fragment (rRF) originating from the 5’-end of 18S rRNA subunit (Fig 4A). While this rRF is not classified as a miRNA in mirBase, recent findings in Zebrafish show that it may interact with Argonaute proteins in a miRNA-like matter [5, 25]. Low detection of this rRF in regular 5P-seq (Fig 4B) would have been overlooked in other studies. Nonetheless, unlike 5X-type libraries that showed very similar rRF profiles, 5P-type libraries were at first dissimilar (Fig 4A-B). A more detailed view on the rRNA subunits revealed, however, that most peaks were present in both the 5P-tag and regular 5P libraries but varied in detection level (Fig 4C-E). Together this indicates that the 5’ RNA ligation strategy—by either using long or short oligos/adapters—severely affects the enrichment of specific rRF, which could make cross-study validation challenging.

We also noticed a 5’ bias in 5P-type and a 3’ bias in 5X-type libraries (Fig 4A-B), and therefore classified each fragment by their 5’ start or 3’ end of a mature rRNA subunit. This confirmed that rRFs generated from the 5’ terminals were enriched in 5P-seq, while rRFs deriving from the 3’ terminals were enriched in 5X-seq (Fig 4F). Importantly, 5XP-seq contained rRFs originating from both the 5’ and 3’ terminal but kept them separated in the 5P-tag and 5X-notag sub-libraries, respectively. These differences indicate that 5P-type libraries lose sRNA primarily at the 3’ end, possibly as a consequence of interference with RNA-modifications during reverse transcription. Interestingly, 5’ rRF bias has recently been reported in humans using an independent 5P-seq kit {Nätt, 2020 #2760;Hua, 2019 #2565}.

### Coverage of small tRNA fragments strongly depends on library preparation strategy

Like rRFs, tRNA derived sRNA (tRFs) can also originate from the 5’ or 3’ terminal. In addition, internal (i’) fragments, which neither start nor end in the terminals of full-length tRNA have been described [26, 27]. We used these classifications as an independent confirmation of the differences in 5’ and 3’ affinities between the 5P- and 5X-type libraries that we observed in sRNA derived from rRNA. As expected, 5P-seq had a clear 5’ preference, while 5X-seq progressively increased towards the 3’ terminal (Fig 5A). Similar to the rRFs, 5XP-seq contained tRFs originating from both the 5’ and 3’ terminal and successfully separated them into the two sub-libraries.

**Fig 5.**
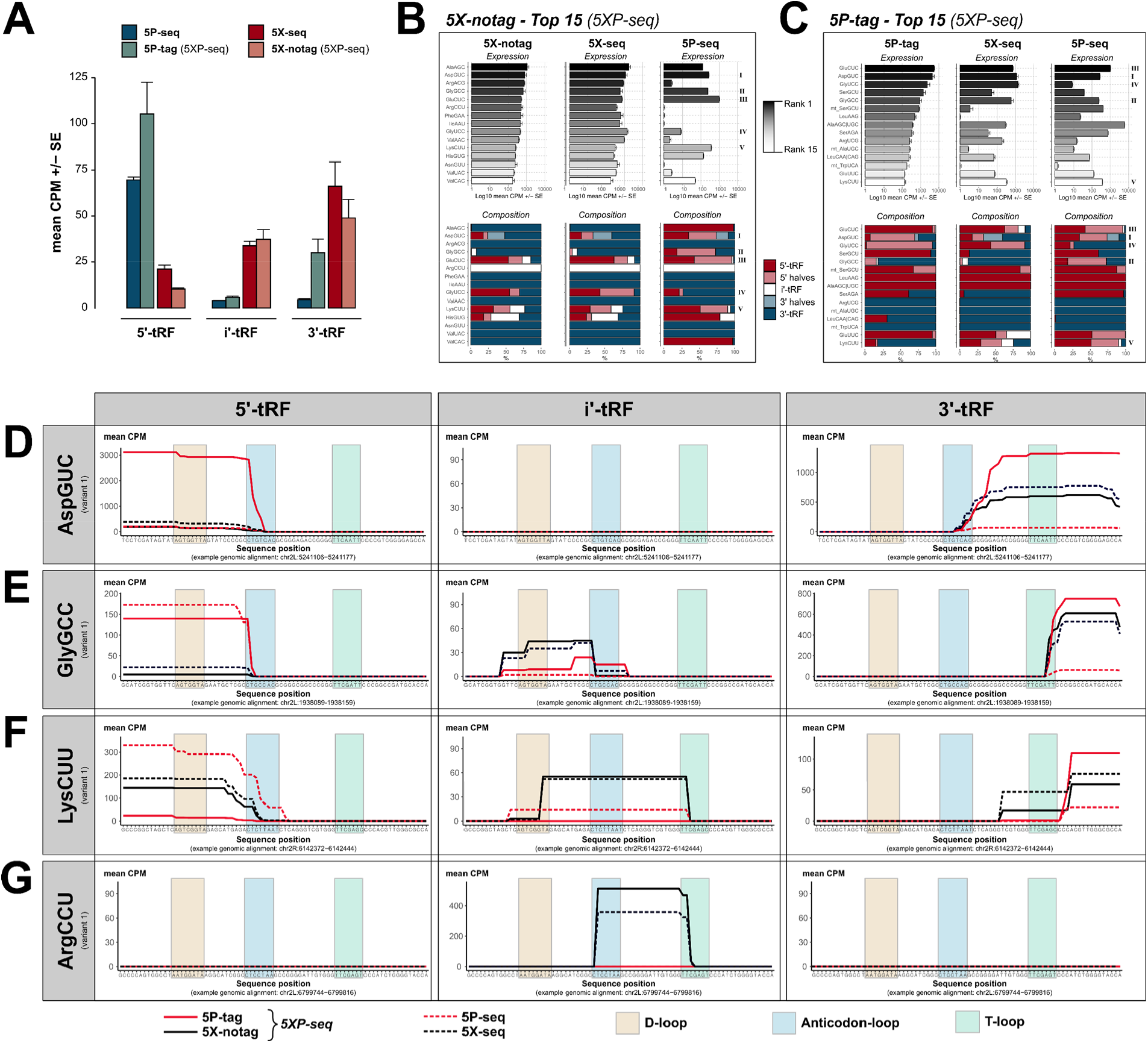
tRNA isodecoder analysis reveals the sRNA complexity generated by different library preparation strategies. Graphs show tRNA derived sRNA (tRFs) grouped by their classification into 5’, i’ and 3’ sub-species, and tRNA isodecoder family. **(A)** Bars show 5’-tRF enrichment in 5P-type libraries and 3’-tRF enrichment in 5X-type libraries. **(B)** Top 15 expressed isodecoders in 5X-notag and their corresponding expression levels (black-grey-white bars) and 5’/i’/3’ ratios (blue/light blue stacked bars) in regular 5X-seq and 5P-seq libraries. **(C)** Same as (B), but the top expressed isodecoders in 5P-tag sub-libraries. **(D-G)** Show tRF coverage plots of a selection of isodecoders, separated by 5’, i’ and 3’ tRF classification (left, middle and right panels, respectively). **(D)** AspGCU, illustrating anticodon-loop cleavage with both 5’ and 3’ halves. **(E)** GlyGCC, illustrating a 5’ half tRF cleaved in the anticodon-loop and a short 3’ tRF cleaved in the T-loop. **(F)** LysCUU, illustrating many 5’- and 3’-tRFs, but also an i’-tRF resulting from D- and T-loop cleavage. **(G)** ArgCCU showing a i’-tRF cleaved between the anticodon- and T-loops. CPM = counts per million reads.

In addition to 5’, i’ and 3’ stratification, we classified each tRF according to the isodecoder of the mature tRNA in order to better understand the possible biases in the tRF coverage introduced by different library strategies. This enables an overview of tRF subtypes, such as tRNA derived stress-induced small RNA (tiRNA) that are generated by angiogenin dependent anticodon cleavage of specific isodecoders resulting in tRNA halves [28], as well as the recently discovered nuclear internal T-loop tRFs (nitRNA) [7]. As expected, 5X-notag sub-libraries generated primarily i’ and 3’ fragments that was strongly replicated by regular 5X-seq, but poorly replicated by 5P-seq (Fig 5B). Again, while the most covered tRFs in the 5P-tag sub-libraries of 5XP-seq showed similarities to regular 5P-seq, we found more variability on the isodecoder level (Fig 5C). In highly abundant isodecoders, there were also an enrichment of sRNA species derived from the 5’ end of tRNAs but this was less pronounced in less abundant isodecoders.

In line with the rRNA analysis, more in-depth analysis of a selection of tRNA isodecoders showed that most tRFs were detected by each method but detection levels varied greatly across methods. For example, the 5P-tag sub-library was superior in detecting both the 5’and 3’ halves generated by angiogenin-mediated cleavage of AspGUC, which generates a known tiRNA (Fig 5D). In another confirmed tiRNA originating from GlyGCC, the 5’-half was detected in both 5P-tag and regular 5P libraries (Fig 5E). At the 3’-end a short fragment generated by T-loop cleavage were detected in the 5P-tag, 5X-notag and regular 5X libraries but to a much lesser degree in regular 5P libraries (Fig 5E). LysCUU has previously been shown to generate many different tRFs [7, 29]. This was confirmed by 5X-notag, regular 5X and regular 5P, but to a lesser degree in 5P-tag libraries (Fig 5F). Interestingly, primarily 5X-notag and regular 5X libraries detected a weakly expressed long T-loop internal tRFs comparable to the nitRNA that was recently detected in human sperm [7]. Similarly, 5X-notag and regular 5X, but not 5P-tag and regular 5P, detected a short nitRNA in ArgCCU generated by cleavage in the T- and anticodon-loops that was almost identical to the one previously reported in human sperm [7].

Together these findings show that the choice between 5P-sensitive and 5P-insensitive methods are critical when studying tRFs using sequencing. Thus, using a method that conserves the diversity of both phosphorylated and non-phosphorylated sRNA—like 5XP sRNA-seq—is highly desirable.

## DISCUSSION

Here we presented a novel method—5XP sRNA-seq—that sequences sRNA with and without 5’ phosphorylation (5’-P) using the same library. Our protocol was optimized for low starting material which makes it attractive for use with precious samples. By ligating a short 5’-P oligonucleotide tag, we demonstrated that 5’-P sRNAs can be separated from other sRNAs in the downstream analysis. We reported multiple examples in which a one-sided approach would have had severe consequences for the interpretation and cross-validation of sRNA experiments. We also provided evidence that ligating a short oligonucleotide tag enhances the recovery of 5’-P sRNAs, such as piRNA and miRNA.

It has become increasingly clear that the 5’ terminal of RNA is subject to diverse modifications. As this is the start site for transcription, it means that if the first molecule is a nucleotide, it will initially have a triphosphate. If transcribed by RNA polymerase II, to become for example mRNA, a cap consisting of modified guanine (G) nucleotide is first added to the initial nucleotide during transcription. Primary piRNAs are also transcribed by RNA polymerase II and receive a similar cap. These 5’ caps regulate the stability and determine the downstream fate of the transcript. The process of generating sRNAs from longer precursors by nuclease digestion will result in either -P or -OH terminals, depending on the nuclease. Intriguingly, it was recently shown that sRNAs are capped as well, indicating that not all caps are added during transcription but rather after the cleavage to shorter transcripts [13, 30]. In addition to the guanine-based caps, recent discoveries show that the sRNA may have other types of 5’-caps.

First reported in bacteria, and suggested to be an alternative to the eukaryotic m^7^G cap, 5’ NAD caps have now been demonstrated in several species including humans (see review: [31]). Interestingly, NAD^+^, as well as NADH and dpCoA, can be incorporated into RNA during transcription initiation, thus serving as non-canonical initiating nucleotides [32]. In human cells, the amount of RNA with 5’-NAD caps change in response to shifting cellular NAD concentrations [33], while data from Arabidopsis shows enrichment of NAD-caps for transcripts involved in redox responses [31]. These two findings, together with the fact that NAD+/NADH is one of the most important intracellular redox pairs, suggests that NAD-capping of RNA may regulate gene expression as a function of the cell’s redox state. Moreover, members of the NUDIX hydrolase superfamily have been shown to remove not only 5’-NAD, but also 5’-FAD and 5’-CoA, indicating that metabolite-containing 5’ caps might be a more widespread phenomenon than previously thought [34]. Whether NAD or other metabolic 5’-caps are used to regulate sRNA, however, is yet to be discovered.

These recent findings demonstrate the urge to develop new sequencing methods that not only include 5’ modifications other than phosphate, but also correctly identify each modification in downstream analyses. By using RNA ligases with specificity toward different 5’ terminals together with unique oligonucleotide tags, 5’XP sRNA-seq is a first step towards a more holistic sequencing approach. By tagging 5’-P in this way, our data suggest that 5’-P sRNAs constitute only 10-20% of the total pool of sRNA in drosophila embryos. Therefore, up to 90% of sRNA species may have previously escaped our attention since most commercial kits are based on 5’-P dependent adapter ligation. Using 5’XP sRNA-seq on RNA from *Drosophila* embryos, we discovered several new sRNA species without 5’-P, including cross-species validation of a zebrafish 5’-rRF from the 18S rRNA subunit [25], and nitRNA derived from internal tRNA T-loop cleavage, previously described only in human sperm [7].

## CONCLUSION

By combining 5’ insensitive library strategies with 5’ sensitive tagging, we have demonstrated an innovative strategy for solving an intrinsic bias in modern sRNA sequencing. Our results represent an important step towards a new generation in sRNA sequencing that can explore the complete world of sRNAs in single low-input experiments. Future technologies will be aimed at further expanding the number of specific terminal RNA modifications that can be identified by genome-wide sequencing approaches. We anticipate discovering a rich repertoire of functional RNA modifications that will greatly expand our understanding of how genomes are regulated.

## Methods

### Experimental Methods

#### Reagents and Oligos

Many of the reagents used in 5XP-seq are included in the NEBNext Multiplex Small RNA Library prep kit for Illumina (New England Biolabs), and are referred to as ‘NEB-kit’ below. Since we used lower input (30 ng) than the recommended 100 ng total RNA, we diluted the 3’SR Adapter, 5’SR Adapter, and SR RT Primer 1:2 in nuclease free H2O. The SR Primer and all Index Primers used for amplification, were used in original concentrations.

Custom oligos not included in the NEB-kit were all HPLC-purified: 5P-tag RNA oligo (long: 5’-UGGCAACGAUC-3’; short: 5’-UGGGAUC-3’; both 3.75 μM).

R1R DNA Adapter (5’ Phos-GAT CGT CGG ACT GTA GAA CTC TGA ACG TGT AG-SpcC3 3’; 100 μM). Adenylation of R1R was done using 5’ DNA Adenylation kit (New England Biolabs) according to manufacturer’s instructions and cleaned with Oligo Clean & Concentrator (Zymo Research) as described here [23]. Oligos was eluted and diluted with nuclease-free water to 10 μM.

Illumina multiplex PCR primer (5’ -AAT GAT ACG GCG ACC ACC GAG ATC TAC ACG TTC AGA GTT CTA CAG TCC GAC GAT C-3’; 10 μM)

Since *Drosophila melanogaster* expresses large amounts of 2S rRNA in the same size as many sRNA, we blocked this transcript by adding anti-sense oligos at the 5’ RNA ligation (First), and at the cDNA ligation steps (Second). For species with less 2S rRNA, these blocking oligos can be exchanged for nuclease-free H2O or custom oligos targeting other transcripts that may occupy a large amount of the sequencing capacity.

First 2SrRNA block oligo (5’-TAC AAC CCT CAA CCA TAT GTA GTC CAA GCA-SpcC3 3’; 10 μM) [35].

Second LNA 2SrRNA block oligo (5’-TGC-**T**TG-GAC-**T**AC-ATA-**T**GG-TT**G**-AGG-**G**TT-G**T**A-SpcC3 3’; 10 μM, where bold letter indicate LNA incorporation).

#### RNA Extraction from Drosophila Embryos

RNA was isolated from *Drosophila melanogaster* (W1118) embryos 0.5-2.5h of age. Embryos were decorionated in pools of 50 embryos using sodium hypochlorite 3.5% (RECTAPUR) and washed extensively with RNase free H2O. RNA was isolated using the miRNeasy Micro kit (Qiagen) according to manufacturer’s instructions. In brief, 500 μl Qiazol (Qiagen) was added to the embryos and homogenized for 2 min at 40 Hz using a Tissue Lyser LT (Qiagen) and 5 mm stainless steel beads (Qiagen). Phase separation was done by mixing 100 μl of Chloroform followed by centrifugation at 12000xg for 15 min in 4°C. RNA was then collected by columns, washed and eluted in 14 μl nuclease free H2O. RNA integrity was confirmed by Bioanalyzer (Agilent) and concentrations were determined by Nanodrop (ThermoFisher).

#### 5’XP sRNA-seq Library Preparation

A side-by-side comparison of the workflows of the different methods used in this study is available in Supplementary Fig 1.

The 5’XP sRNA-seq method includes the following steps:

|  |  |  |  |
| --- | --- | --- | --- |
| i. | Ligation of 3' RNA adapter | 18.5 h | Proceed immediately (over-night) |
| ii. | Ligation of 5' RNA tag | 2.5 h | Proceed immediately |
| iii. | cDNA synthesis | 2.5 h | Pause point |
| iv. | Ligation of 5' cDNA adapter | 3.0 h | Pause point |
| v. | Barcoding & Amplification | 1.5 h | Pause point |
| vi. | Size selection | >12 h | Pause point (over-night) |
| vii. | Pooling barcoded samples | 3 h | Pause point |

In detail, the 3’ adapter was first ligated using 0.5 μl of 3’SR Adapter for Illumina (NEB-kit) added to 30 ng of input RNA diluted in 3.0 μl nuclease free H2O. Samples were heated to 70°C for 2 min and immediately cooled on ice. A mix of 5 μl 3’Ligation Reaction Buffer 2x (NEB-kit) and 1.5 μl of 3’Ligation Enzyme Mix (NEB-kit) were added and incubated for 18 h at 16°C over-night. A 2S rRNA blocking oligo was added simultaneous to hybridization of the reverse transcription primer by adding a mix of 0.5 μl of First 2SrRNA block oligo, 0.5 μl of SR RT primer (NEB-kit) and 1.75 μl of nuclease-free water to each sample followed by incubation at 90°C for 30 s, 65°C for 5 min, 37°C for 15 min, and finally 25°C for 15 min. 5’-tagging was then performed by adding 0.5 μl 5P-tag RNA oligo (denatured at 70°C for 2 min), 0.5 μl 5’Ligation Reaction Buffer (NEB-kit) and 1.25 μl of 5’Ligation Enzyme Mix (NEB-kit) followed by a 1 h incubation at 25°C. To synthesis cDNA, a mix of 4 μl First strand Synthesis Reaction Buffer (NEB-kit), 0.5 μl Murine RNase Inhibitor (NEB-kit) and 0.5 μl ProtoScript II RT (NEB-kit) was added to each sample, and incubated first at 50°C for 1 h, and then at 70°C for 15 min. Enzymatic inactivation and hydrolysis of RNA was done by incubating samples at 95°C for 3 min with 17.5 μl nuclease-free water, 10 μl 0.5M EDTA and 2.5 μl 5M NaOH. After room temperature equilibration pH was lowered by adding 1 μl 5M HCl. Samples were then cleaned using Oligo Clean & Concentrator kit (Zymo Research) according to manufacturer’s recommendation but with an extra wash in 250 μl DNA wash buffer (Zymo Research) and cDNA elution in 11 μl of nuclease-free water. Ligation of pre-adenylated R1R adapter with 2S RNA blocking was done by adding 10 μl of cDNA, 2 μl 10x NEBuffer 1 (New England Biolabs), 2 μl 50 mM MnCl2, 2 μl Thermostable 5’App ligase (New England Biolabs), 4 μl 10 μM pre-adenylated R1R Adapter, and 0.5 μl 10 μM Second LNA 2SrRNA block oligo, followed by 2 h incubation at 65°C. Ligated cDNA samples were cleaned and eluted in 24 μl Elution Buffer (Qiagen) using Oligo Clean & Concentrator (Zymo Research) according to manufacturer’s instructions. Clean samples were amplified by adding 25 μl 2x Phusion High-Fidelity PCR master mix (ThermoFisher), 1 μl 10 μM Illumina Multiplex Primer, 1 μl 10 μM Illumina barcode (NEB-kit) to 23 μl of cDNA, and incubated in a thermocycle at 98°C for 5 s, followed by 15 cycles of 98°C for 5 s, 60°C for 10 sec, 72°C for 30s, ending with 72°C for 1 min. Amplified libraries were cleaned using Agencourt AMPure XP (Beckman Coulter), and size selected for 130-200 nt fragments on a pre-casted 6% polyacrylamide Novex TBE gel (Invitrogen). Gel extraction was done using Gel breaker tubes (IST Engineering Inc) in DNA Gel Elution Buffer (NEB-kit). Disintegrated gels were incubated at 37°C for 1h on a shaker, quickly frozen for 15 min at −80°C, followed by another incubation for 1 h. Remaining gel debris were removed by Spin-X 0.45 μm centrifuge tubes. The libraries were precipitated overnight at −80°C by adding 1 μl of GlycoBlue (Invitrogen) as co-precipitant, 0.1 times the volume of Acetate 3M (pH 5.5), and three times the volume of 100% ethanol. Library concentrations were estimated using QuantiFluor ONE ds DNAsystem on a Quantus fluorometer (Promega). Pooled libraries were sequenced on NextSeq 500 with NextSeq 500/550 High Output Kit v2.5, 75 cycles (Illumina).

#### 5’P sRNA-seq Library Preparation

Library preparation was done with NEBNext Multiplex Small RNA Library prep kit for Illumina (New England Biolabs) according to the manufactures instructions except for downscaling all samples to half volume and using 30 ng of input RNA (100 ng recommended) with an appropriate 1:2 dilution of all adapters. This corresponds to the 5XP-seq preparation as above, but with the following changes:

In Step [ii.], 5’ RNA ligation was performed using 0.5 μl 5’ SR Adapater (NEB-kit) instead of the 5P RNA oligo.

In Steps [iii-iv.], clean up of cDNA, ligation of the 5’ DNA adapter and the Second LNA 2SrRNA oligo block was not performed. The cDNA reaction mix was immediately amplified according to kit protocol.

In Step [v.], 20 μl cDNA reaction mix was amplified by adding 25 μl of Long Amp Taq 2x Master Mix (NEB-kit), 1.25 μl SR primer for Illumina (NEB-kit), 2.5 μl nuclease-free H_2_O, 1.25 μl 10 μM Illumina barcode (NEB-kit) and incubated, starting with 94°C for 30 s, followed by 15 cycles of 94°C for 15 sec, 62°C for 30 sec, 70°C for 15 sec, ending with 70°C for 2 min.

#### 5’X sRNA-seq Library Preparation

5X-seq library preparation was identical to 5XP-seq library preparation, except for the following adjustments:

In Steps [ii-iii.], 5’ RNA ligation of the 5P-tag RNA oligo and the First 2SrRNA block was not performed. The cDNA reaction mix was therefore compensated by adding 0.5 μl 5’Ligation Reaction Buffer 10x (NEB-kit) and 1.75 μl nuclease-free H_2_O.

## Computational Methods

### Quality Control and Pre-processing

Raw fastq data files have been deposited in Sequence Read Archive under the accession number PRJNA658107. All libraries passed Illumina’s default quality control. Demultiplexed fastq files were downloaded from BaseSpace using BaseMount (Illumina) and lanes were merged for each sample. For all library-types the 3’ adapters were trimmed using Cutadapt 2.3 [36] with following input: -a AGATCGGAAGAGCACACGTCTGAACTCCAGTCACAT --discard-untrimmed -- nextseq-trim=20 -O 5 -m 5. This specifically trims nextseq type sequences between 5-70 nt with at least 5 nt of the 3’ adapter present in the 3’ end and discarding all other sequences. For 5XP-seq libraries we reloaded the trimmed sequences into Cutadapt using the following input: -g TGGCAACGATC -m 5 --untrimmed-output. This saves sequences with or without the 5’-tag in separate fastq output files. All trimmed fastq files were quality filtered using fastq_quality_filter -q 20 -p 80 -v available in the FASTX Toolkit 0.0.14 (https://github.com/agordon/fastx_toolkit), which only retain sequences with PHRED score > 20 in more than 80% of nucleotides. The integrity of trimmed and quality filtered fastq files were further verified using FastQC 0.11.9 (https://github.com/s-andrews/FastQC).

### Sequence Counting, Filtering and Annotation

Data was summarized over counts of unique sequences across all samples. Tools for such sequence-based approaches are available—such as Sports and MintMap [26, 37]—and may be contrasted against feature-based counting approaches, where sequences are counted over genomic features—such as the genomic coordinates of miRNA and piRNA. One benefit of unique sequence-based approaches is that the sequence of the original read is maintained during counting, which is often lost du initial mismatch allowance in feature-based approaches. Here we counted unique sequences in trimmed fastq files using customized scripts in R 3.4.4 [38]. For parallel and efficient large data processing, foreach 1.5.0 [39] and the data.table [40] packages were used. Reading and processing fastq-files were done using the ShortRead package [41]. Prior to normalization, noise was reduced by only including sequences with at least 5 counts in 100% of samples within a given method (5P-seq, 5X-seq, 5XP-seq etc). This dataset contained 330 971 unique small RNA sequences with mean total counts per sample of 8 993 458. Normalized counts in reads per million (rpm) were generated by dividing the individual sequence counts (reads) with the total sequence counts for each sample. For some analysis that targeted highly abundant sRNAs we further reduced the data by only including sequences with at least 10 rpm in 100% of the samples within a given method included in the analysis. Some critical values of the datasets are available in Table 1.

Sequence annotation was performed by mapping unique sequences against small RNA sequence reference databases using bowtie 1.2.2 [42]. This was done in cycles, where each cycle allowed one additional mismatch, from 0 to 3 mismatches. A sequence was only destined for another bowtie annotation cycle if it failed to align to any of the databases in the previous cycle. Fasta reference sequence files were attained from the following databases: miRNA = miRbase and Ensembl, tRNA = GtRNAdb (nuclear) and Ensembl (mitochondrial), rRNA = Ensembl (both nuclear and mitochondrial), piRNA = pirBase, Other sncRNA = Ensembl. We used dm6 versions across all databases. To resolve multimapping issues when sequences matched multiple databases within the same mismatch category, we applied the following hierarchy miRNA > Mt_tRNA > tRNA > MT_rRNA > rRNA > piRNA > other sncRNA. A sequence mapped to miRNA with 1 mismatch and piRNA with 0 mismatch, would therefore annotate as piRNA, while if both had 0 mismatch it would annotate as miRNA.

### Statistical Methods and Visualization

All statistical analysis was done in R 3.4.4 [38]. Data visualization was primarily done using the ggplot2 package in R [43] and finalized using Inkscape 0.92. For fitting the saturation plots we used non-linear least square regression with an asymptotic self-starter (r functions: nls and SSasymp). For hierarchical clustering and correlation plots we used the corrplot package [44]. For tRNA/rRNA coverage we used custom scripts wrapped around the vmatchPattern function in the Biostrings package [45]. A version of this script has been published [7] and are available here: https://github.com/Danis102/Natt_et_al_2019_Human_Sperm_Rapid_Response_to_Diet/blob/master/S1_Text.R. Reference sequences were obtained from GtRNAdb (tRNA) and Ensembl (rRNA). GtRNAdb ss-files were used to map tRNA loops. Future scripts for 5’XP sRNA-seq will be posted here: https://github.com/Danis102/.

## ACKNOWLEDGEMENTS

The study was supported by grants from The Swedish Research Council (201503141; https://www.vr.se/english.html; received by AÖ), Knut and Alice Wallenberg Foundation (Wallenberg Academy Fellow, 2015.0165; https://kaw.wallenberg.org/wallenberg-academy-fellows; received by AÖ), Ragnar Söderberg (Fellow in Medicine 2015; https://ragnarsoderbergsstiftelse.se; received by AÖ). The funders had no role in study design, data collection and analysis, decision to publish, or preparation of the manuscript.

