## Supplementary Figures for "5’XP sRNA-seq: Efficient Identification of Transcripts With and Without 5’ Phosphorylation Reveals Evolutionary Conserved Small RNA"

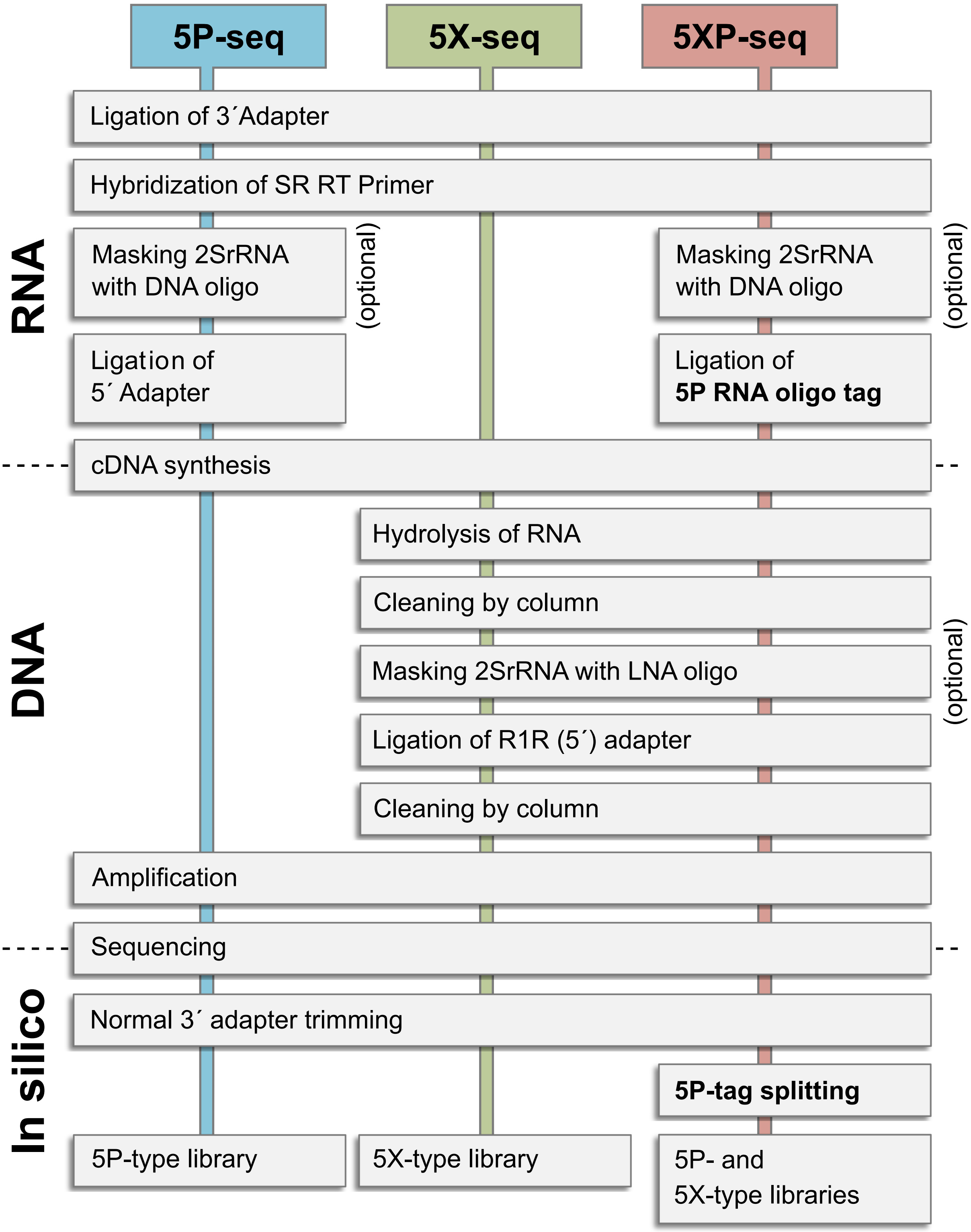


**Supplementary Fig 1. Flow chart comparison between the 5P-, 5X- and 5XP-seq methods.** In order to cover more types of small RNA we combined the 5P-seq with 5X-seq. Regular 5P-seq first ligates the 3´ adapter and then the 5´ adapter to RNA prior to cDNA synthesis. 5X ligates only the 3´ adapter to RNA while the 5´ adapter (R1R) is ligated after cDNA synthesis. 5XP-seq generates 5P- and 5X-type libraries from the same sample. Instead of ligating a 5´ adapter to the RNA, in 5XP-seq ligates a short oligo tag that later identifies 5´ phosphorylated sRNA using the 5X-seq workflow.

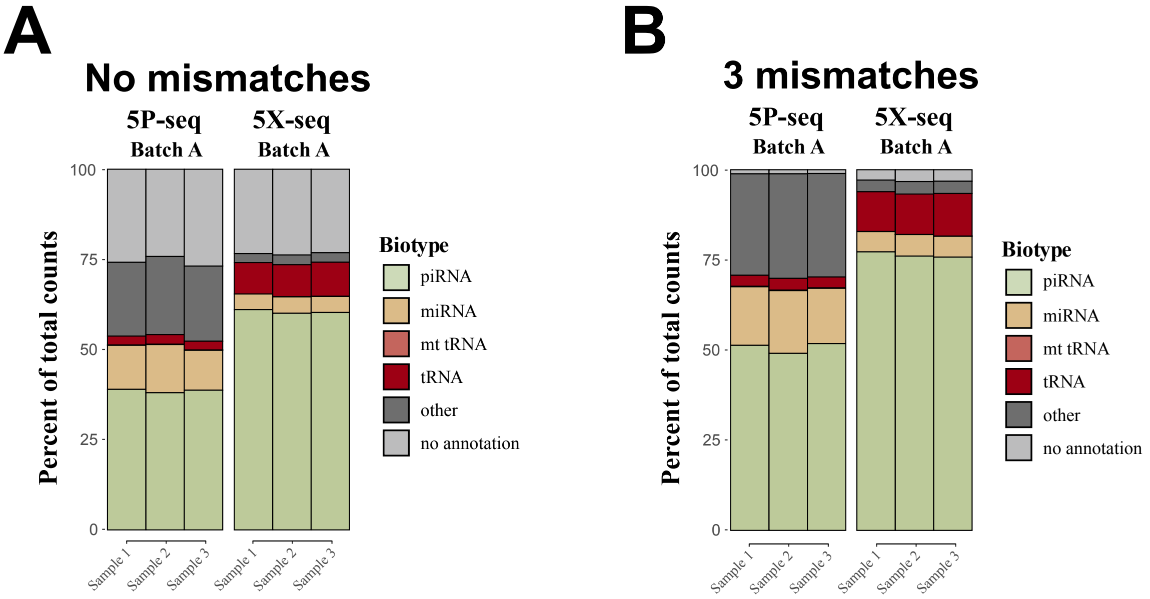


**Supplementary Fig 2. Biotype composition in 5P and 5X-seq libraries without rRNA.** Stacked bars with the biotype compositions of 5P and 5X libraries from Batch A isolated from the same pool of RNA from drosophila embryos. Identical to Fig 2, except that the small RNA annotating to rRNA and Mt_rRNA was removed. Only unique sequences that passed 10 CPM in all samples of a method (either 5P or 5X) was included. **(A)** Shows the result when allowing 0 mismatches in the mapping between read sequence and reference sequence, while **(B)** shows the result when 3 mismatches was allowed. mt=mitochondrial. CPM=Counts per million reads.

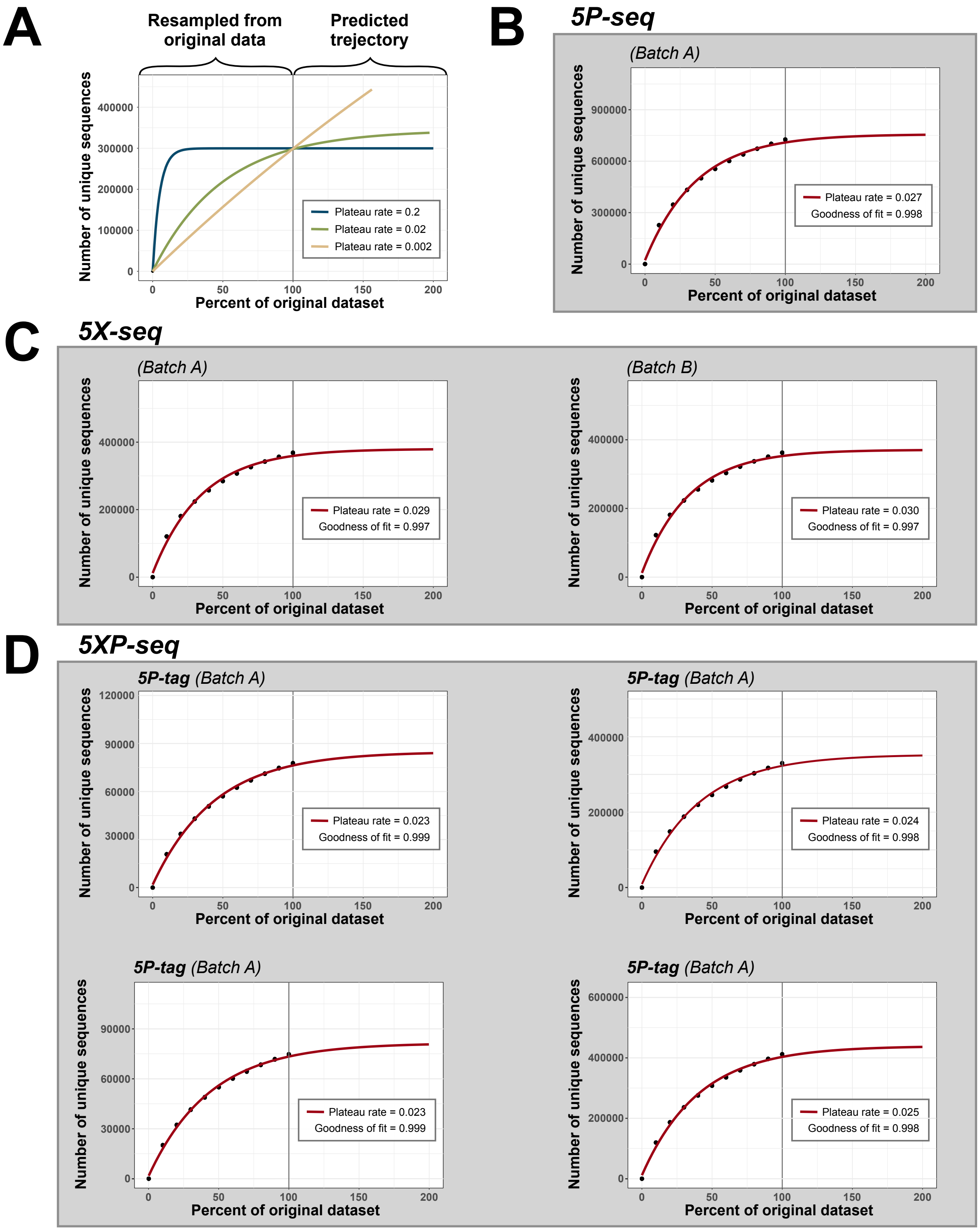


**Supplementary Fig 3. The 5P-tag sub-libraries of 5XP-seq need less unique sequences to reach a plateau.** Saturation plots with asymptotic non-linear regression estimating the percent of library needed to reach a theoretical plateau where no new sequences will be found no matter how much more sequence is aquired. **(A)** Shows the principle of the saturation analysis, where counts of unique sequences are resampled 10 times at different ratios (0,10... 80, 90%) from the original counts (100%). Asymptotic cruve is then fitted to the data to predict the curve trajectory at >100% and the rate constant--Plateau rate--is extracted. Colors indicate model behavior at different Plateau rates, where blue shows oversaturated, yellow undersaturated and green a typical library. **(B)** Shows the saturation plot for regular 5P-seq where black dots are the 10 resamples. **(C)** Shows the same for the two batches of regular 5X-seq, while **(D)** shows the same for the different batches and sub-libraries of 5XP-seq. Note that the Plateau rate only differs slightly between libraries but the 5P-tag sub-libraries need fewer unique sequences to reach a plateau.
